# Multitaxon assessment reveals inconsistent biodiversity responses to forest structural complexity in temperate forests

**DOI:** 10.1101/2025.10.28.685007

**Authors:** Cesare Pacioni, Kris Verheyen, An Martel, Lander Baeten, Frank Pasmans, Bram Catfolis, Tosca Vanroy, Leni Lammens, Luc Lens, Diederik Strubbe

## Abstract

Increasing forest structural complexity is a key objective of future-proof forest management, with potential benefits for biodiversity. However, empirical evidence for consistent biodiversity-structure relationships across taxa is still limited. We investigated whether structurally more complex forests support greater species richness and higher multidiversity across taxonomic and functional groups in 19 mature forest plots in Flanders, Belgium. As one of the most densely populated and urbanized regions in Europe, with limited and highly fragmented forest cover, Flanders provides a particularly informative and policy-relevant context to test structure–biodiversity relationships. Its forests, often dominated by a few tree species and subject to long-term anthropogenic pressures and management, represent a realistic gradient of structural complexity. Structural complexity was quantified using a Structural Complexity Index (SCI), and biodiversity was assessed using a multidiversity index integrating scaled species richness across five taxonomic and seven arthropod functional groups. Using mixed-effects models and multivariate Bayesian analyses, we tested both direct effects of SCI on biodiversity and the covariation in species richness among groups. Contrary to expectations, SCI was not a consistent predictor of multidiversity, and most groups showed weak or inconsistent responses. These findings might suggest that structural complexity alone may be insufficient to enhance biodiversity in simplified forests.

**Highlights:**

- Forest structural complexity does not consistently predict biodiversity.
- Responses vary among taxonomic and functional groups.
- Cross-taxon congruence in diversity is limited and scale-dependent.
- Keywords: Forest structural complexity; Multidiversity, Species richness; Functional groups; Sustainable forest management

## 1. Introduction

Forests cover nearly 40% of Europe’s land and support much of the continent’s terrestrial biodiversity, providing essential ecosystem services (State of Europe’s Forests, 2020). However, centuries of intensive land use and the widespread application of conventional forest management practices, particularly those favoring even-aged monocultures, have led to the simplification of forest structure and the homogenization of forest composition across many of Europe’s forested landscapes (Zerbe, 2023). This reduction in heterogeneity in stand age, vertical layering, canopy cover, deadwood availability, and tree species composition (Schall & Ammer, 2013; Siitonen et al., 2000; Wirth et al., 2009) has significant ecological consequences, as many forest-dwelling species rely on structural diversity for habitat, foraging, and reproduction. For example, specialist species, such as cavity-nesting birds and saproxylic insects, are particularly sensitive to reductions in deadwood or old-growth features (Bujoczek et al., 2020; Kajtoch et al., 2013; Lassauce et al., 2011; Vítková et al., 2018). Recent ecoacoustic research in temperate European forests shows that bird species richness is significantly higher in stands with greater structural heterogeneity, especially in forests with high tree species richness, tree size variability, and snag height variability, highlighting the importance of maintaining diverse forest structures to support avian diversity (Shaw et al., 2024). Given the ecological implications of structural simplification, there has been a growing interest in prioritizing structurally complex, mixed-species forests with uneven-aged canopies (Messier et al., 2015), strongly endorsed by the EU’s Nature Restoration Regulation. This shift is reflected in recent forest management approaches across Europe aiming to enhance biodiversity and increase ecosystem resilience to climate extremes, pest outbreaks, and anthropogenic disturbances (Fahey et al., 2018; Muys et al., 2022). These more complex stands tend to have a wider range of microhabitats (Paillet et al., 2017), which can increase niche availability, shelter opportunities, and food resources for a wide range of species (Tews et al., 2004; Toivonen et al., 2023; Wagenaar et al., 2025). Consequently, increasing and maintaining higher levels of forest structural complexity is increasingly viewed as a priority in sustainable, future-proof forest management, with potential co-benefits for biodiversity conservation and ecosystem functioning (Bauhus et al., 2009; Dove & Keeton, 2015; Keeton, 2006).

The relationship between forest structural characteristics and biodiversity has long been a central theme in ecological research. However, findings are often context-dependent, varying across taxonomic groups, spatial scales, and forest types (Bohn & Huth, 2017; Müller et al., 2023; Oettel & Lapin, 2021; Schall et al., 2018; Storch et al., 2023). Structural indicators such as vertical stratification, tree size diversity, and deadwood volume are commonly used because they can be easily quantified and are closely linked to habitat quality, making them suitable proxies for biodiversity in forest inventories (Sabatini et al., 2015; Storch et al., 2018). National forest monitoring programs (e.g., the German Federal Forest Inventory (BWI) and the Flemish Forest Inventory, Vlaamse Biosinventaris) routinely assess these attributes (Vidal et al., 2016a; Vidal et al., 2016b). Although an increasing number of studies support a positive relationship between structural complexity and overall biodiversity, most of this research focuses on individual taxonomic groups. Enhanced structural complexity has been shown to promote beetle diversity and biomass through the presence of multiple forest elements (Rappa et al., 2022), increase fungal species richness via retention of deadwood and aboveground biomass (Dove & Keeton, 2015), and support higher diversity of drosophilid communities in multilayered temperate forests compared to structurally simpler boreal ones (Tanabe et al., 2001). However, focusing on a single taxon may obscure broader ecosystem-level patterns and limit the generalizability of results. In addition, many studies conflate structural complexity with tree species diversity, making it challenging to isolate the independent contribution of forest structure to biodiversity. For example, Juchheim et al. (2020) showed that structural complexity tends to increase with tree species richness, particularly in mixed forests with broadleaved species, thereby complicating efforts to separate structural and compositional effects. By contrast, Springer et al. (2024) and Storch et al. (2023) found that structural variables such as mean tree diameter can predict biodiversity even in species-poor stands. This suggests that structural complexity may have an independent effect on biodiversity, beyond its association with tree species richness. However, these effects were inconsistent across groups (Wagenaar et al., 2025), underscoring the need to distinguish the influence of forest structure from that of species composition. Clarifying this relationship is especially important in forests dominated by a single tree species, where compositional diversity is inherently low but structural variation may still play a role in shaping biodiversity patterns.

As a more integrative approach to assessing ecosystem-level biodiversity responses, Allan et al. (2014) introduced the concept of multidiversity, a composite metric that captures the average proportional richness across multiple taxonomic groups. To avoid dominance by taxa with inherently high species counts, species richness within each group is first standardized to its observed range (scaled between 0 and 1), after which these values are averaged across all groups, giving equal weight to each. By summarizing biodiversity in this balanced way, multidiversity can reveal emergent patterns that are often missed by single-taxon approaches, offering a more ecologically meaningful picture of community-level responses to environmental variation. Applying this framework to forest ecosystems can help identify whether structurally complex stands support consistently high biodiversity across groups, or whether overall richness masks divergent taxon-specific responses. Such multitaxon approaches are essential because different groups often respond differently to structural features (Blasi et al., 2010; Elek et al., 2018; Lengyel et al., 2016; Oettel & Lapin, 2021; Penone et al., 2019; Rigo et al., 2024). These group-specific patterns can reflect underlying ecological traits, including resource dependencies, trophic roles, and sensitivity to disturbance. Arthropod functional groups are particularly informative in this context, as their contrasting ecological roles lead to distinct responses to forest structural and compositional attributes. For example, detritivore abundance is typically shaped by the amount and quality of dead organic matter, whereas herbivores are more directly influenced by tree species diversity, which determines host plant availability (O’Brien et al., 2017).

Multidiversity metrics help integrate such taxon-specific responses by summarizing average proportional richness across groups, thereby providing an ecosystem-wide indicator of biodiversity. However, while multidiversity provides an integrated measure of biodiversity that gives equal weight to each taxonomic group, it does not reveal whether those groups respond similarly or differently to environmental gradients. A complementary perspective emerges from examining how species richness co-varies across taxonomic and functional groups. Such analyses clarify the extent to which different components of biodiversity co-respond to structural forest attributes and thus allow us to assess the consistency of biodiversity patterns across groups. Understanding these patterns is important for informing forest management strategies. For example, if taxa show similar responses, targeted structural interventions could yield broad biodiversity benefits. If responses are decoupled, a more tailored, group-specific approach may be required. Thus, examining covariation adds an important dimension to biodiversity assessment and supports the development of more nuanced and effective conservation strategies in structurally complex forests.

Here, we examined the relationship between forest structural complexity and biodiversity in mature forest stands in Flanders (northern Belgium), a region that offers a particularly informative context for studying structure–biodiversity relationships. Flanders is one of the most densely populated and urbanized regions in Europe (504 inhabitants/km²), with forest cover limited to approximately 11% and intense competition for land use (Govaere & Leyman, 2022; Statistics Belgium, 2025). Forests are typically small and highly fragmented, embedded in agricultural and urban matrices which together cover over 70% of the landscape. These pressures, combined with atmospheric nitrogen deposition and a long history of intensive land use, have been shown to impair forest functioning, reduce biodiversity, and diminish ecosystem service provision (De Keersmaeker et al., 2015; Decocq et al., 2016). Flemish forests are generally dominated by a set of economically important tree species, including pedunculate oak (*Quercus robur*), European beech (*Fagus sylvatica*), and Canadian poplar (*Populus × canadensis*). While many managers have shifted toward multifunctional management, the forest structural complexity often remains relatively low, offering a narrow but realistic gradient for evaluating the independent effects of forest structure on biodiversity.

We established a network of 19 forest plots between Ghent and Brussels, spanning gradients of structural complexity within stands dominated by the above-mentioned oak, beech, and poplar tree species. Structural complexity was quantified using a Structural Complexity Index (SCI), adapted from the multifunctionality framework of Manning et al. (2018) by Catfolis et al. (2023). Biodiversity was assessed using the multidiversity index developed by Allan et al. (2014), which averages scaled species richness across taxonomic groups to give equal weight to each (see above). A range of taxa representing different ecological functions and trophic levels was systematically sampled across plots. We evaluated the influence of SCI using two complementary multidiversity indices: one averaged across all taxonomic groups, and another calculated for functionally defined groups within the phylum Arthropoda, based on feeding strategies and life-history traits. In addition to these integrative measures, we examined SCI effects separately for each taxonomic and arthropod functional group. Our analyses thus address both the direct effects of structural complexity on group-specific richness and the extent to which diversity patterns co-vary across groups in response to forest structural complexity. We hypothesized, first, that forest plots with higher structural complexity would support greater multidiversity, due to increased ecological heterogeneity and niche availability. Accordingly, ecosystem-level biodiversity, captured by the multidiversity index, is expected to peak in structurally rich stands. Second, we predicted that this relationship would be underpinned by a high degree of covariation among taxonomic and functional groups. That is, species-rich plots for one group (e.g., birds) would also be species-rich for others (e.g., arthropods or mammals), indicating a shared response to structural habitat features. Such congruent patterns would suggest that structural complexity acts as a unifying driver of biodiversity across forest biota.

## 2. Material and Methods

### 2.1. Plot characterization and forest structural complexity

This study is part of the FORESTER research project (Catfolis et al., 2023, 2025; De Schuyter et al., 2025; Lammens et al., 2024; Terriere et al., 2022; Vanroy et al., 2024, 2025). This is a large-scale study in the southern part of Flanders, investigating the effect of forest structural complexity on stress and diseases in small mammals, birds and amphibians. This study was conducted in the same network of forest plots (Figure 1). The 19 one-hectare plots were situated between Ghent and Brussels, on soils ranging from loam to sandy loam, in a region with a mean annual temperature of ∼11D°C and precipitation of 800–900 mm. Plots were selected to represent a gradient in SCI while minimizing variation in forest size, history, and environmental conditions. Forests were grouped by dominant tree species, pedunculate oak (*Quercus robur*), European beech (*Fagus sylvatica*), or Canadian poplar (*Populus × canadensis*), defined as the species covering more than 60% of the canopy. All selected forests are at least 3Dha, date back to before 1850, and are separated by at least 1Dkm. Each forest contained a 1-hectare ‘core plot’, selected to be representative of the forest and located near its center. These core plots were inventoried following Catfolis et al.’s (2023) Structural Complexity Index (SCI) protocol, based on the multifunctionality framework proposed by Manning et al. (2018). This protocol uses a combination of stand-level visual assessments, vegetation surveys, and structural measurements. These included estimates of canopy cover, stand age, number of tree layers, tree species mixture, and cover and composition of tree, shrub, and herb layers. Tree diameter distributions and deadwood metrics were also recorded using concentric sampling circles and line intersect transects. Based on these variables, a SCI was calculated using a multi-component scoring system with predefined sub-indices for forest structure, woody layer, herb layer, and dead wood. These components are reported separately in our results (see Figure 2) to illustrate how total SCI varies across plots and forest types. While the protocol allows analysis of these sub-indices, data analyses (not shown) indicated that they did not show consistent patterns across plots or taxa. Therefore, in this study, we focus exclusively on the total SCI as a summary measure of forest structural complexity. Taxa included in this study were Aves, Mammalia, Arachnida, Syrphidae, Apoidea, and Arthropoda. Although Syrphidae, Apoidea, and Arachnida all belong to the phylum Arthropoda, they were treated as separate groups in this study. This decision was made because the Arthropoda group was assessed at the order level and classified into functional groups based on feeding strategy, while all other taxa were identified to the species level. Moreover, the Arthropoda dataset originated from a distinct sampling protocol targeting functional diversity within orders, which overlapped with taxonomic groups already represented. To avoid redundancy and inconsistencies due to differences in taxonomic resolution and sampling methods, the Arthropoda data were analyzed independently. Arthropod orders were classified into functional groups based on their feeding habits (Lu et al., 2024; Tobisch et al., 2023). The functional groups considered were detritivores, herbivores, non-feeders, omnivores, parasites, pollinators, and predators.

**Figure 1.**
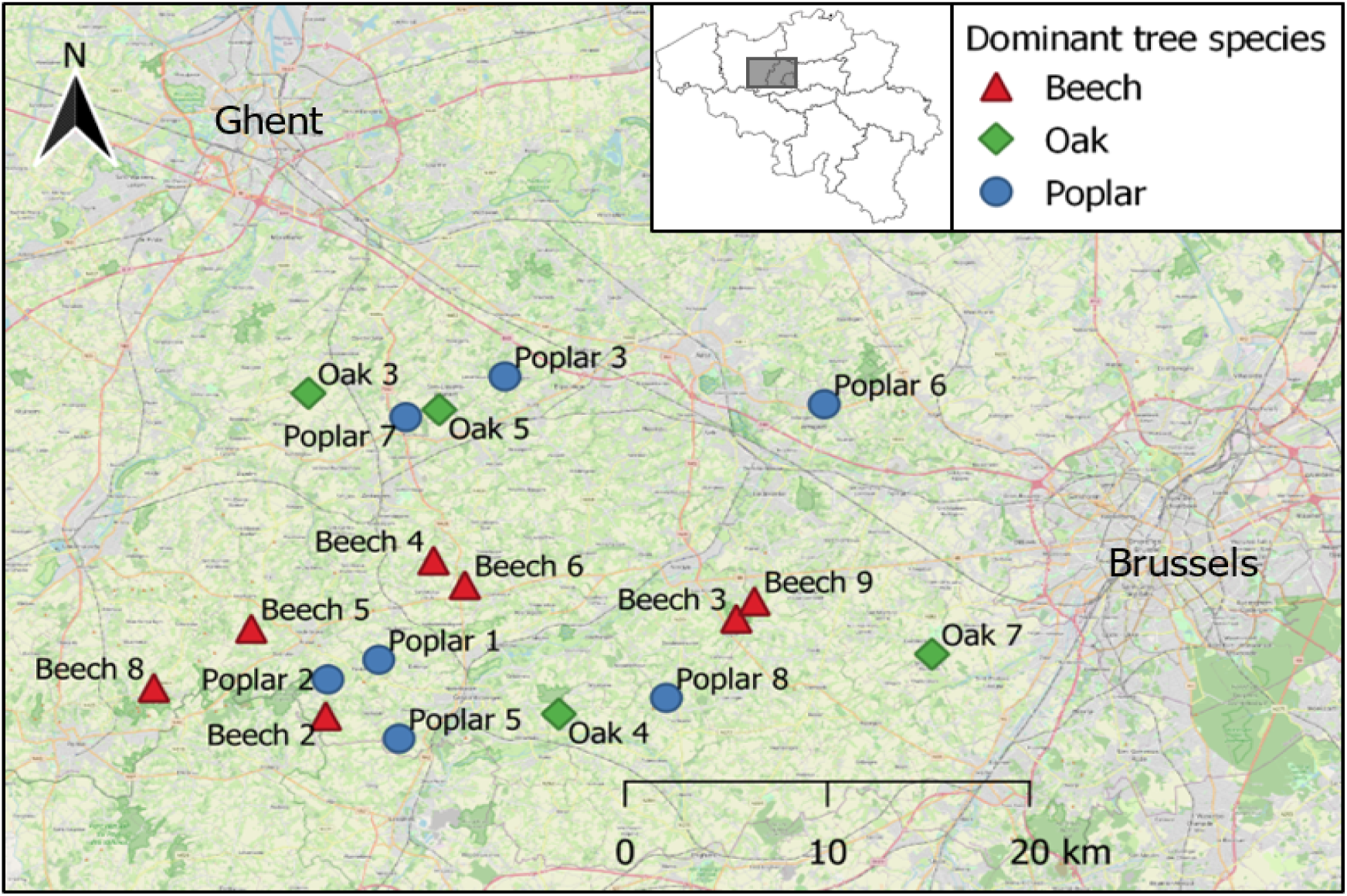
Map of the study sites in Flanders (Belgium). The colored symbols show the dominant tree species of that forest. From Catfolis et al. (2023) and Vanroy et al. (2025).

**Figure 2:**
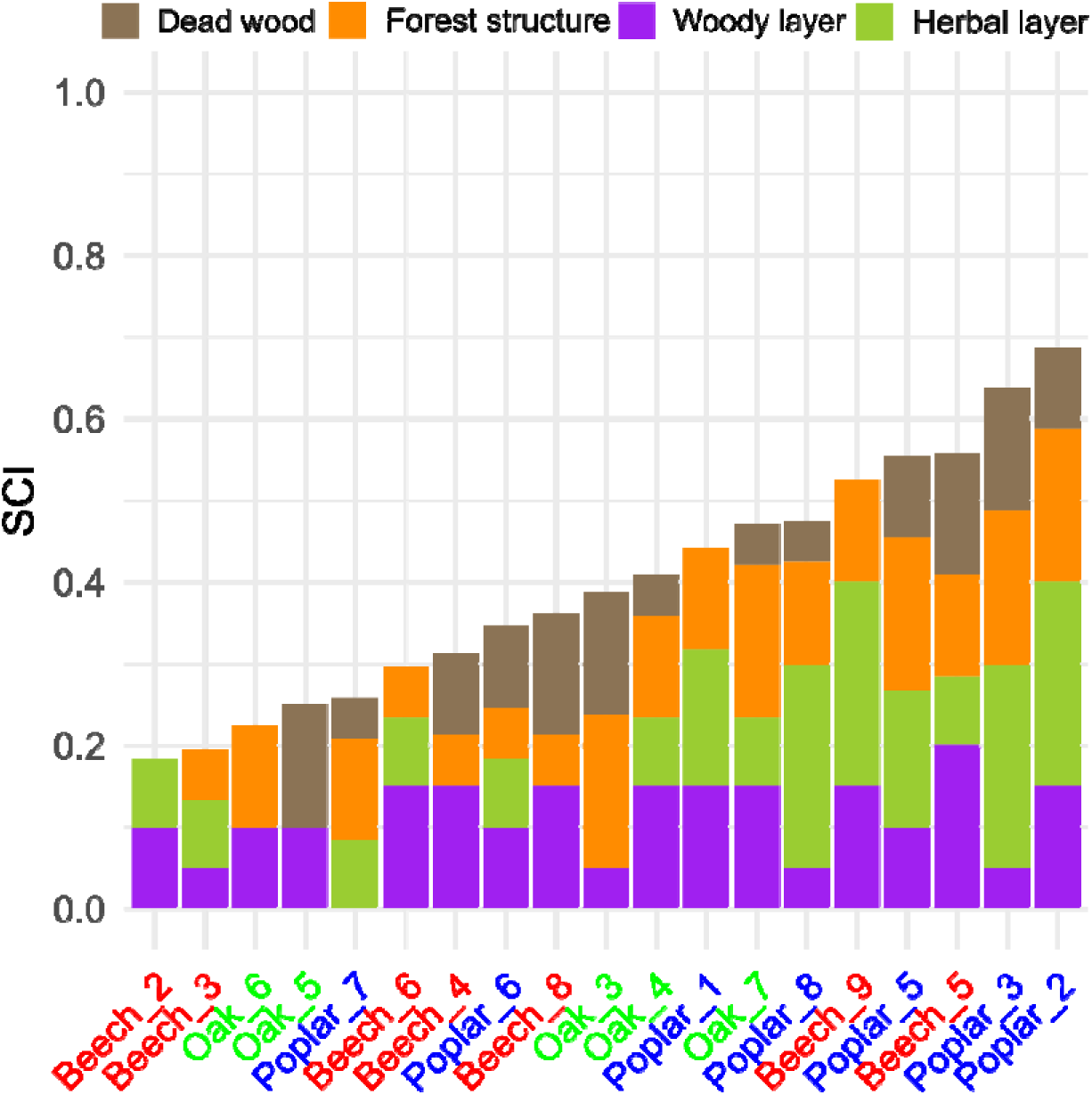
Gradient in forest structural complexity across 19 study plots. Each stacked bar represents the Structural Complexity Index (SCI) for a forest plot, decomposed into its four components: forest structure (orange), woody layer (purple), herbal layer (olive green), and dead wood (brown). Forest plots are ordered by total SCI value from lowest to highest, highlighting variation in structural complexity across beech (red), oak (bright green), and poplar (blue) stands.

### 2.2. Sampling

This study integrates data on six taxonomic groups collected following established protocols. For detailed methodologies, see the respective cited publications. Bird sampling was conducted between February and April 2021 (Catfolis et al., 2023). A total of 712 individuals (17 species) were captured using 12m wide mist nets (Ecotone, mesh size 16×16 mm) deployed at study plots. Captured birds were individually sexed, aged by plumage (Demongin, 2016), and fitted with a unique metal ring from the Belgian Ringing scheme. Regarding mammals, rodents were trapped in mid-August to mid-September 2021 following Vanroy et al. (2025). A grid of 49 live traps (Trip Trap with wooden extensions) baited with oats, raisins, peanut butter, and mealworms was deployed in a 7×7 layout with 5 m spacing. Traps were pre-baited one night before sampling, set one hour before sundown, checked after 3 hours, and closed after an additional 3 hours. Captured individuals, mainly wood mice (*Apodemus sylvaticus*) and bank voles (*Myodes glareolus*), were identified to species and sex. A total of 314 were captured across the study plots, comprising 123 *A. sylvaticus* and 187 *M. glareolus*. Wild pollinator communities (Syrphidae and Apoidea) and spiders (Arachnida) were sampled using UV-colored pan traps (yellow, white, blue) mounted 1 m above ground on wooden platforms, avoiding interference with bramble vegetation (De Schuyter et al., 2025; Smedt et al., 2023). Two trap setups were placed per 1-ha core plot, spaced by at least 20 m. Pan traps were filled with water and biodegradable soap and equipped with drainage holes to prevent overflow. Sampling occurred throughout 2021, with contents collected every two weeks from April 21 to June 7, and every four weeks from July 8 to October 4 (totaling eight collections and 166 sampling days). Samples were sieved (1 mm mesh) and preserved in 70% ethanol before identification to species level. Arthropods were sampled biweekly from July to October 2023 within 1-hectare core plots (Catfolis et al., 2025). The late-season sampling period was selected to ensure more stable weather conditions and reduce the impact of short-term fluctuations in temperature and precipitation, which are increasingly common in spring due to climate change (Forrest, 2016; Kharouba et al., 2018; van Dijk et al., 2024). This approach enhances comparability across sites by minimizing interannual variability in arthropod emergence and activity. Despite the timing, key taxa such as Coleoptera, Araneae, Isopoda, and Hymenoptera remain abundant and ecologically relevant during late summer and early autumn in temperate forests (Crowley et al., 2023; Isaia et al., 2015). To further control for weather-related variation, sampling was restricted to calm, dry days. Arthropods were collected using three complementary methods targeting different vertical strata: pitfall traps for the litter layer, sweep netting for the herbaceous layer, and the beating sheet method for the shrub layer. Each plot was sampled nine times using sweep netting and beating sheets, and eight times using pitfall traps.

### 2.3. Statistical analysis

To evaluate the effect of SCI on overall biodiversity, multidiversity was calculated at two levels: (i) averaged across all taxonomic groups, and (ii) across functionally defined groups within the phylum Arthropoda. For both levels, linear mixed-effects models were fitted using the lme4 package (Bates et al., 2015) in R, with mean multidiversity as the dependent variable. SCI, dominant tree species, and forest area were included as fixed effects together with all interactions. Models included taxonomical or functional group identity as a random intercept to account for group-level differences in baseline multidiversity. Mean multidiversity index was calculated by averaging the scaled multidiversity values across all taxonomic groups within each plot. To further investigate the influence of SCI on taxonomic multidiversity, a threshold-based analytical approach was applied (Allan et al., 2014). Species richness values were first standardized within each taxonomic group by scaling them between 0 and 1 across all plots, such that 1 corresponded to the highest observed richness for that group and 0 to the lowest. For each plot, a threshold-based multidiversity index was then calculated by counting the number of taxonomic groups with scaled richness values exceeding a given threshold. This procedure was repeated using three different thresholds (0.25, 0.5, and 0.75) to assess the sensitivity of the results to the threshold parameter. The resulting index, ranging from 0 to 5, was used as the response variable in generalized linear models (GLMs) with a Poisson error distribution. Predictor variables included the SCI, tree identity, and plot area. Model overdispersion was evaluated using the dispersion ratio, calculated as the Pearson residual deviance divided by the residual degrees of freedom. In cases where overdispersion was detected (i.e., dispersion ratio > 1.5), the models were refitted using a negative binomial distribution.

To assess group-specific responses, univariate linear models were fitted separately for each major taxonomic and functional group. The response variable was the group-specific scaled species richness, with SCI, dominant tree species, forest area, and their interactions included as fixed effects. For each univariate model, the overall effect of the categorical predictor dominant tree species was assessed using an F-test. To evaluate differences among individual tree species (beech, oak, and poplar), pairwise comparisons of estimated marginal means (EMMs) were conducted with Tukey adjustment for multiple testing. These post-hoc comparisons were performed using the emmeans package (Lenth et al., 2023). All models assumed Gaussian error distributions and, where appropriate, were fitted using restricted maximum likelihood (REML). Forest area was log-transformed to reduce skewness and then scaled (standardized to mean 0 and SD 1) to improve model fitting and interpretation. Statistical significance was evaluated at a threshold of p ≤ 0.05. Model assumptions of normality and homoscedasticity were checked via residual diagnostics.

To complement the multidiversity analyses, we also examined how SCI influences covariation in species richness among groups. To quantify this, we employed a Bayesian multivariate modelling framework using the *brms* package (Bürkner, 2017) in R. Separate models were constructed to assess covariation across taxonomic groups and across functionally defined arthropod groups. In each model, species richness in the included groups was modelled jointly as a multivariate response, with SCI, dominant tree species, and forest area included as predictors of both the mean richness and the residual correlations (i.e. covariation) among groups. Models were fitted using Hamiltonian Monte Carlo sampling across four independent chains, each run for 20,000 iterations with a warm-up phase of 4,000 iterations. The target acceptance probability was increased (adapt_delta = 0.95) to improve sampling stability and ensure convergence. Model performance and predictive accuracy were evaluated using approximate leave-one-out cross-validation based on Pareto-smoothed importance sampling. Differences in expected log predictive density were used to compare relative support among candidate models. To examine patterns of residual covariation among the response variables, the residual correlation matrix was extracted from the posterior distribution of each model. These residual correlations were summarized as posterior means and 95% credible intervals, and used to assess whether diversity among different taxa or functional groups co-varied independently of the modeled environmental drivers. Correlations with credible intervals excluding zero were interpreted as evidence of non-random residual co-variation. All statistical analyses were performed using R software v. 4.2.2 (R Core Team 2022), details of which are available in the RMarkdown supplementary file.

## 3. Results

### 3.1. Multidiversity analyses

First, we tested whether structurally complex forests support higher overall biodiversity (multidiversity across all taxonomic groups), but no statistical support was found for this hypothesis. Multidiversity was significantly lower in oak stands compared to beech (Estimate = –0.93, p = 0.038), and a strong positive interaction between forest area and oak also emerged (Estimate = 4.94, p = 0.003). The three-way interaction between SCI, oak, and forest area was significant and negative (Estimate = –13.29, p = 0.010), indicating that this positive oak–area effect weakened when structural complexity was high. No other terms were statistically significant (all p > 0.10). Second, we evaluated effects of SCI on biodiversity across arthropod functional groups. This model showed no significant main effect of SCI on multidiversity (Estimate = –0.48, p = 0.202).

Across all thresholds tested, the Poisson GLMs indicated no statistically significant effects of SCI, dominant tree species, or forest area on the number of taxonomic groups whose scaled richness exceeded the threshold, i.e. those with relatively high richness values. At the lowest threshold (0.25), most plots contained multiple groups with richness values above the threshold, resulting in low variability in the multidiversity index (range: 3–5). SCI had a small, non-significant positive effect (Estimate = 0.22, p = 0.80). At the intermediate threshold (0.5), slightly more variation in the threshold-based multidiversity index was observed (range approximately 2–5), but the effect of SCI remained non-significant (Estimate = –0.83, p = 0.48). The model using the highest threshold (0.75) yielded a more pronounced negative effect of SCI (Estimate = –3.24), though this, too, did not reach statistical significance (p = 0.21).

### 3.2. Univariate taxon and arthropod functional group specific analyses

Univariate, taxon-specific models revealed heterogeneous responses of individual taxonomic groups to forest structural complexity (SCI) and covariates (Figure 3). Only for Aves, SCI had a significant (and negative) effect on scaled species richness index (Estimate = –1.69, *p* = 0.037). Forest area showed no significant effect in the presence of interactions (*p* = 0.24). The model for arthropods showed that species richness was higher in poplar stands compared to beech, but only as a significant interaction with area (Estimate = 5.06, *p* = 0.030; model R² = 0.40). Among the arthropod groups assessed separately, Syrphidae species richness exhibited no significant main effects, but showed marginal interactions involving SCI and area (Estimate = 6.95, *p* = 0.060) and a marginal negative three-way interaction with poplar (Estimate = –9.50, *p* = 0.060; model R² = 0.49). Apoidea species richness was significantly lower in poplar stands compared to beech (*p* = 0.038), while the negative effect of SCI was marginal (*p* = 0.073; R² = 0.45). For Mammalia and Arachnida, none of the predictors nor interactions had a significant effect on scaled species richness (*p* > 0.29 for all terms). Univariate, arthropod functional group-specific models showed mixed and often non-significant effects of SCI on multidiversity (Figure 4). A significant negative effect of SCI was found for pollinators (Estimate = –2.37, p = 0.017), while no other group exhibited a significant SCI effect. Detritivores and omnivores showed significant or marginally significant three-way interactions involving SCI, tree identity, and area. Parasites had a marginally non-significant negative effect of SCI (p = 0.078). Herbivores, non-feeders, and predators displayed low explanatory power (R² < 0.57, adjusted R² often negative), and none of the predictors were significant.

**Figure 3.**
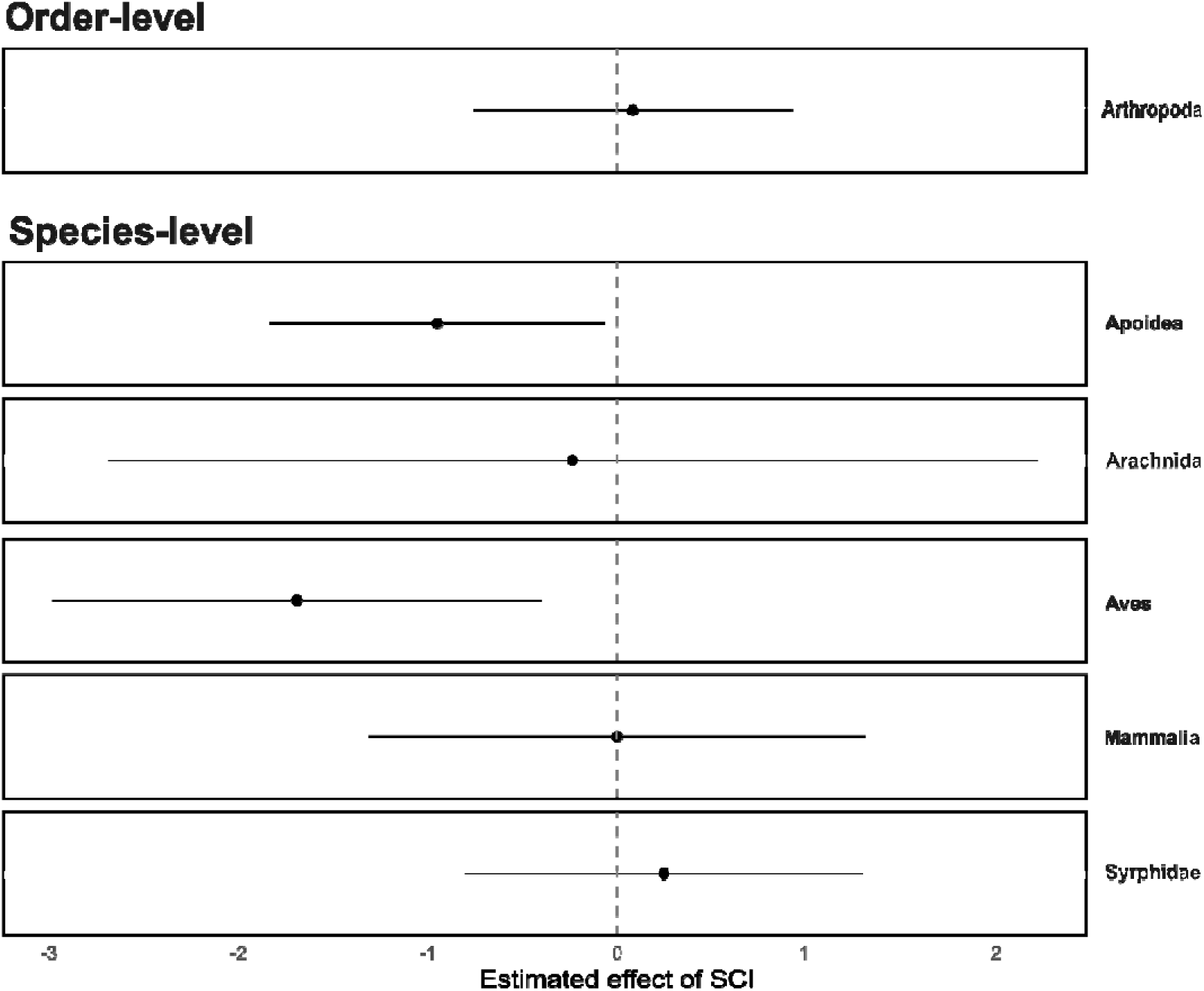
Estimated effects (±95% CI) of forest structural complexity (SCI) on taxon-specific multidiversity. Estimates are based on separate linear models for each taxonomic group, controlling for dominant tree species and forest area.

**Figure 4.**
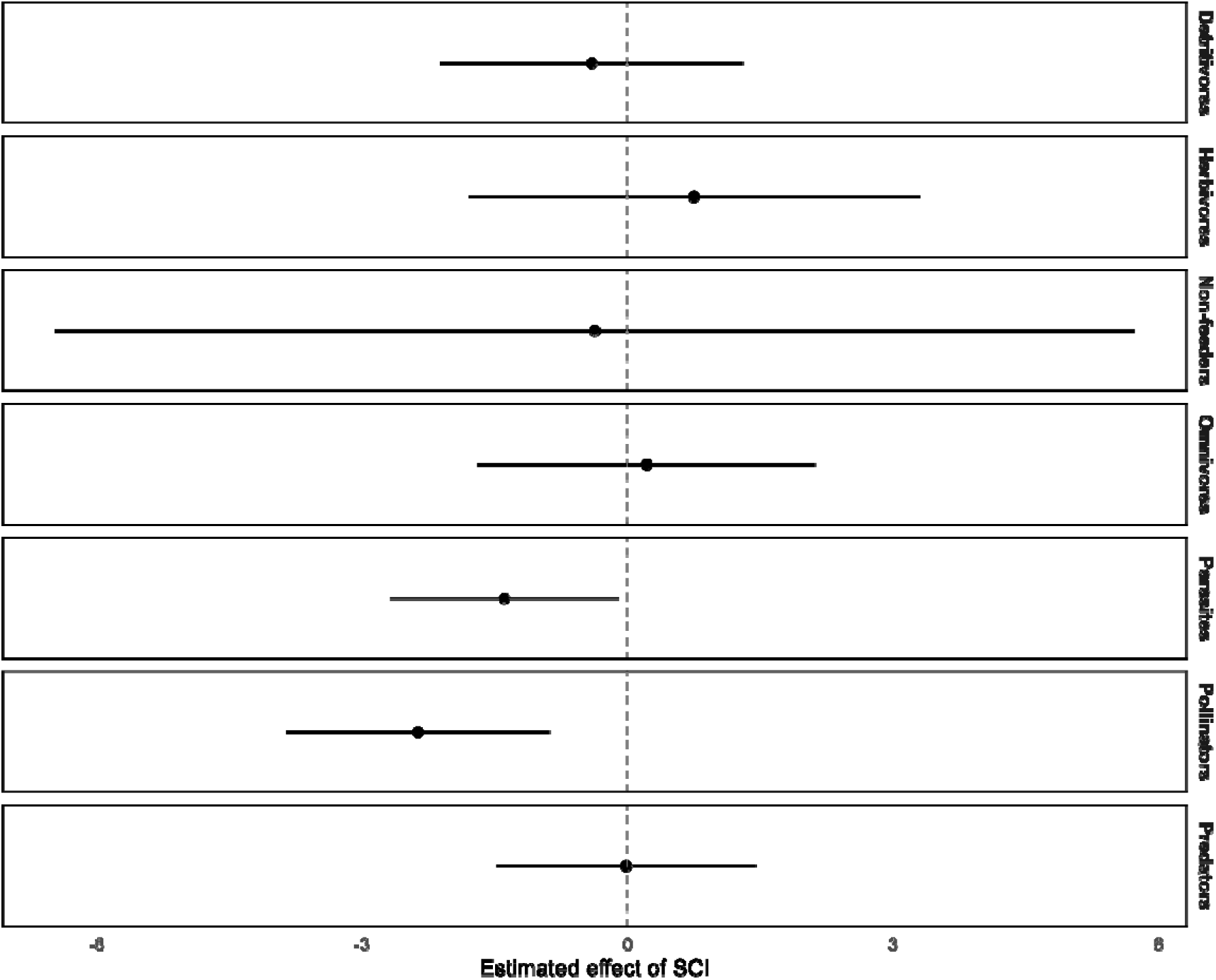
Estimated effects (±95% CI) of forest structural complexity (SCI) on arthropod functional groups. Estimates are based on separate linear models for each functional group, controlling for dominant tree species and forest area.

### 3.3. Covariation analyses

Across taxonomic groups, we found no credible evidence for a correlation in species richness (Table S1), nor any indication that SCI, dominant tree species, or forest area influenced richness covariation among groups (i.e. the null model provided the best fit, see RMarkdown supplementary file). Consequently, residual correlations between taxonomic groups were generally low to moderate. Apoidea and arthropod multidiversity were negatively correlated (mean = –0.48, 95% CI: –0.76 to – 0.09), whereas Arachnida and Apoidea multidiversity showed a positive association (mean = 0.41, 95% CI: 0.00 to 0.72). Also across arthropod functional groups, a null model fit best (RMarkdown supplementary file), and correlation and residual correlations among arthropod functional group multidiversity values were generally low (Table S2). Across all models, most residual correlations between pairs of multidiversity values had 95% credible intervals overlapping zero (RMarkdown supplementary file).

## 4. Discussion

Contrary to our initial hypotheses, the results provide limited support for a positive effect of forest structural complexity on species richness across taxonomic groups and arthropod functional groups. Species richness did not increase significantly with higher values of the Structural Complexity Index (SCI), and multidiversity was not consistently associated with structural gradients. While certain patterns emerged, these effects were not uniform across groups. Univariate analyses showed heterogeneous and often weak to non-existent responses to SCI. Species richness did not covary strongly among taxonomic or arthropod functional groups, and SCI did not emerge as a reliable integrative proxy for biodiversity under the conditions studied.

### 4.1. Taxon- and group-specific responses to forest structure are weak and inconsistent

Although structurally complex forests are generally presumed to offer more ecological niches and microhabitats, thereby supporting a greater number of species and trophic diversity (Bazzaz, 1975; Jung et al., 2012; St. Rose & Naithani, 2025; Taboada et al., 2010; Wagenaar et al., 2025), our findings did not reveal a consistent or strong positive response between structural complexity and species richness across taxonomic or arthropod functional groups. One likely reason for the weak and inconsistent biodiversity–structure relationships is the relatively narrow (though realistic) gradient in structural complexity present in our study system (Figure 2). This limited range largely reflects the deliberate choice to control for confounding variables such as soil type, minimum forest age, and patch size, which tend to reduce environmental noise. Although plots were selected across three dominant tree species (i.e., oak, beech, and poplar), the overall structural variation in Flemish forests remains modest compared to more heterogeneous or old-growth systems. To capture a broader and potentially nonlinear biodiversity–structure relationship, future research could include both larger and smaller forest fragments, younger or coniferous stands, or truly old-growth forests, thereby expanding the SCI gradient. Additionally, incorporating sites with greater variation in management intensity or disturbance regimes may further increase structural complexity variation within the regional context, improving the ability to detect stronger biodiversity responses. Such sampling could, for example, indicate that biodiversity–structure relationships may be nonlinear, with marked increases in diversity only occurring beyond certain structural thresholds that were not captured in our sampling. If that is the case, our results may underestimate the potential of SCI to reflect biodiversity patterns in forests with greater structural differentiation, particularly those with older, more diverse tree communities. Supporting this hypothesis, Evans et al. (2017) demonstrated that species richness of ectomycorrhizal fungi, epiphytic lichens, and ground flora shows sharp threshold responses along a gradient of basal area in dieback-affected beech forests. These results highlight that biodiversity responses can remain relatively flat across much of a structural gradient, then shift abruptly near compositional or structural tipping points. Likewise, Estavillo et al. (2013) found that small mammal assemblages exhibited taxon-specific threshold responses to forest cover loss, with forest specialists collapsing below 30% cover and generalists peaking in diversity near this threshold due to increased landscape heterogeneity. Together, these studies suggest that even modest structural simplification, if crossing critical thresholds, can abruptly alter community composition and functional diversity. Although these studies address different spatial scales (stand vs. landscape), both studies converge on the idea that biodiversity–structure relationships may be nonlinear and group-dependent, shaped by species’ habitat specialization and dispersal ability.

The weak and inconsistent biodiversity responses observed in our plots is also in line with the hypothesis that increasing structural complexity within species-poor stands may offer limited biodiversity benefits (Felton et al., 2016). While structural enrichment in these forests (e.g., through thinning, deadwood retention, or understory development) may increase habitat heterogeneity to some extent (Asbeck et al., 2023), the absence of tree species-level compositional diversity may constrain the types of structures available and limit the variety of niches supported. In other words, even structurally enriched stands dominated by a single species may lack the functional and architectural diversity required to sustain high biodiversity. This aligns with recent findings in German forests, which showed that both carbon storage and multidiversity were highest in structurally complex stands dominated by multiple broadleaved species and where trees were large and presumably older (Springer et al., 2024). In contrast, mostly monoculture conifer-dominated stands, particularly those dominated by pine, consistently supported lower biodiversity and carbon values (Springer et al., 2024).

These findings suggest that high structural complexity, along with its ecological benefits, may be difficult to achieve in largely monospecific stands, but rather requires mixed-species assemblages that contribute diverse traits and habitat features. Supporting this, Juchheim et al. (2020) quantified the relationship between tree species mixing and stand structural complexity using terrestrial laser scanning data from Central European forests. They found an increasing but saturating relationship between species diversity and structural complexity, with the presence of broadleaved trees notably enhancing the structural complexity of conifer-dominated stands. This indicates that converting monospecific stands into mixed forests, particularly by including species with complementary physiological and morphological traits, can promote the development of more complex and functionally diverse forest structures. Such compositional diversity thus appears essential for fostering the structural heterogeneity that underpins richer biodiversity.

### 4.2. Low cross-taxon congruence challenges structural complexity as a biodiversity indicator

Equally important is the observation that cross-taxon congruence in species richness was generally low across plots. This challenges a common assumption in conservation biology, that the richness of one taxonomic or functional group can reliably serve as a surrogate for others (Rodrigues & Brooks, 2007; Westgate et al., 2014). Despite moderate to high residual variation within each group, SCI did not emerge as a consistent driver of covariation in richness across taxa or arthropod functional groups. This suggests that different taxonomic groups and arthropod functional groups may respond to distinct ecological filters or habitat features, even within the same forest stands. While multidiversity aims to integrate such responses into a single metric, our findings caution against assuming that shared environmental drivers (e.g., structural complexity) consistently generate coordinated biodiversity patterns.

The divergent responses of individual taxonomic and arthropod functional groups further illustrate the limitations of applying a single structural index to predict biodiversity patterns across diverse communities. However, even when considering SCI sub-indices targeting the woody layer, herb layer, or deadwood, no consistent patterns emerged across taxa, highlighting that these structural components alone are insufficient to predict biodiversity. Certain bird species and flower-visiting insects might depend on specific microhabitat conditions, such as patchiness of vegetation or availability of particular floral resources, which are not captured at the plot-level by SCI or its sub-indices. Others, such as detritivores and parasites, showed complex interactive effects involving tree identity and area, further emphasizing the importance of species-specific ecological traits and resource requirements. These patterns underscore that the scale at which structural complexity operates differs across taxa. Birds may respond to canopy openness and large-scale heterogeneity, whereas many arthropods depend on fine-scale microhabitats or resource distributions not reflected in plot-level SCI values. Such mismatch in spatial scales between structural metrics and organismal perception may explain why neither SCI nor its sub-indices were not a reliable predictors of cross-taxon patterns in our study. In this context, the scale of analysis has often been shown to affect cross-taxon congruence (Banks-Leite et al., 2011; Dolph et al., 2011; Paavola et al., 2006; Pearson & Carroll, 1999) due to the scale-dependence of biological responses to environmental variation (Infante et al., 2009; Qian & Kissling, 2010; Rooney & Azeria, 2015).

### 4.3. Biodiversity in simplified forests depends on more than structural complexity

Our findings offer limited evidence for a consistent positive relationship between forest structural complexity and species richness. Contrary to our expectations, SCI did not consistently predict biodiversity across taxonomic and arthropod functional groups. Responses were generally weak, varied among groups, and were sometimes even negative. The low cross-taxon congruence further challenges the assumption that one group can serve as an effective surrogate for others. Within this study system and the observed range of structural complexity, SCI was not a consistent predictor of biodiversity patterns. This aligns with a recent meta-review, which found that while certain forest attributes (e.g., deadwood, old trees, and tree cavities) consistently enhance biodiversity, relationships between stand-level structural features and biodiversity are often inconsistent across taxa (Wagenaar et al., 2025). Moreover, the SCI captures the overall structural characteristics of the forest biotope, such as canopy layers, tree density, and deadwood abundance, but it does not directly measure the specific microhabitats, food resources, or environmental conditions that individual species rely on. In a multitaxa context, a single composite index cannot fully represent the diverse habitat requirements of birds, insects, fungi, and other groups, which likely contributes to the heterogeneous and sometimes contrasting responses we observed across taxa.

### 4.4. Study limitations and recommendations for future research

While this study offers valuable insights into how biodiversity responds to forest structural complexity, several limitations should be acknowledged. First, the sampling methods may not have fully captured species richness across all taxa. For example, bird diversity was assessed using mist netting during late winter and early spring, a method known to under-sample canopy-dwelling species and to be affected by weather conditions. Similarly, small mammal sampling was based on a single 7×7 trap grid used over a limited time window, which may have reduced statistical power and led to underestimates of species richness. Second, the study was conducted over a single season and year, which limits the ability to detect seasonal or interannual variation in species occurrence and activity. Finally, the structural complexity gradient in the study area was relatively narrow. Although ecologically realistic, this limited range may have constrained the detection of stronger or nonlinear biodiversity responses. To address these limitations, future studies should consider refining sampling approaches. For bird communities, combining mist netting with complementary methods, such as point counts or acoustic monitoring, could improve species detectability and reduce bias. For small mammals, using multiple trapping sessions and applying capture–recapture or occupancy modelling techniques would offer more robust diversity estimates by accounting for detection probability. Expanding temporal coverage to include multiple seasons and years would allow researchers to capture dynamic changes in biodiversity and better understand how species respond to forest structure over time. Furthermore, studies conducted across a broader gradient of structural complexity, especially including older, less managed, and more compositionally diverse forests, may reveal stronger or more complex biodiversity responses. Finally, future work should integrate additional habitat variables, species-specific ecological traits, and regional context to clarify under which conditions structural indicators effectively reflect biodiversity. Developing such a nuanced, taxon-specific understanding will be important for informing forest management strategies aimed at conserving biodiversity. Beyond these methodological considerations, future work should also explore alternative indicators of biodiversity itself. Our analyses relied on a multidiversity index based on total species richness as the main biodiversity indicator. While this provides an integrated measure across groups, future studies may benefit from focusing on ecologically more specialized subsets of taxa (e.g., saproxylic insects, stenotopic forest species), which may exhibit stronger and more consistent correlations with structural complexity and its components.

## Funding

This study was supported by the UGent GOA project “Forest biodiversity and multifunctionality drive chronic stress-mediated dynamics in pathogen reservoirs (FORESTER)” (grant no. BOF20/GOA/009).

## Data and code availability

All datasets and R script used in this study are available as supplementary material and have been archived on Figshare at https://doi.org/10.6084/m9.figshare.30464372

## CRediT authorship contribution statement

**Cesare Pacioni:** Formal analysis, Visualization, Writing – original draft. **Kris Verheyen:** Conceptualization, Funding acquisition, Supervision, Writing – review & editing. **An Martel:** Conceptualization, Funding acquisition, Supervision, Writing – review & editing. **Lander Baeten:** Conceptualization, Funding acquisition, Supervision, Writing – review & editing. **Frank Pasmans:** Conceptualization, Funding acquisition, Supervision, Writing – review & editing. **Bram Catfolis:** Data curation, Methodology, Investigation, Writing – review & editing. **Tosca Vanroy:** Data curation, Methodology, Investigation, Writing – review & editing. **Leni Lammens:** Data curation, Methodology, Investigation, Writing – review & editing. **Luc Lens:** Conceptualization, Funding acquisition, Supervision, Writing – review & editing. **Diederik Strubbe:** Conceptualization, Methodology, Supervision, Writing – review & editing.

## Declaration of Competing Interest

The authors declare no competing interests.

## Notes

https://doi.org/10.6084/m9.figshare.30464372

### Competing Interest Statement

The authors have declared no competing interest.

