## Supplemental Data 1 for "Multitaxon assessment reveals inconsistent biodiversity responses to forest structural complexity in temperate forests"

**Table S1** Posterior means and 95% credible intervals of residual correlations between taxonomic groups. Correlations are derived from multivariate Bayesian models using different combinations of forest structural variables (SCI, dominant tree species, and forest area). Values are based on the residual correlation matrix estimated in *brms*.

| **Model** | **Correlation** | **Estimate** | **CI Lower** | **CI Upper** | **Includes zero** |
| --- | --- | --- | --- | --- | --- |
| Null | Arachnida – Apoidea | 0.39 | -0.04 | 0.72 | Yes |
| Null | Arachnida – Syrphidae | -0.08 | -0.50 | 0.35 | Yes |
| Null | Aves – Arachnida | 0.00 | -0.43 | 0.42 | Yes |
| Null | Aves – Apoidea | 0.28 | -0.15 | 0.64 | Yes |
| Null | Aves – Mammalia | -0.32 | -0.68 | 0.11 | Yes |
| Null | Aves – Syrphidae | -0.37 | -0.71 | 0.06 | Yes |
| Null | Mammalia – Arachnida | -0.24 | -0.62 | 0.20 | Yes |
| Null | Mammalia – Apoidea | 0.04 | -0.39 | 0.46 | Yes |
| Null | Mammalia – Syrphidae | 0.10 | -0.34 | 0.52 | Yes |
| Null | Syrphidae – Apoidea | 0.07 | -0.36 | 0.48 | Yes |
| SCI + Tree + SCI:log(Area) | Arachnida – Apoidea | 0.22 | -0.29 | 0.65 | Yes |
| SCI + Tree + SCI:log(Area) | Arachnida – Syrphidae | -0.22 | -0.64 | 0.30 | Yes |
| SCI + Tree + SCI:log(Area) | Aves – Arachnida | -0.15 | -0.59 | 0.35 | Yes |
| SCI + Tree + SCI:log(Area) | Aves – Apoidea | -0.29 | -0.69 | 0.22 | Yes |
| SCI + Tree + SCI:log(Area) | Aves – Mammalia | -0.22 | -0.65 | 0.28 | Yes |
| SCI + Tree + SCI:log(Area) | Aves – Syrphidae | -0.42 | -0.77 | 0.07 | Yes |
| SCI + Tree + SCI:log(Area) | Mammalia – Arachnida | -0.21 | -0.65 | 0.30 | Yes |
| SCI + Tree + SCI:log(Area) | Mammalia – Apoidea | 0.28 | -0.23 | 0.69 | Yes |
| SCI + Tree + SCI:log(Area) | Mammalia – Syrphidae | 0.17 | -0.34 | 0.61 | Yes |
| SCI + Tree + SCI:log(Area) | Syrphidae – Apoidea | -0.18 | -0.62 | 0.31 | Yes |
| SCI + Tree + log(Area) | Arachnida – Apoidea | 0.31 | -0.19 | 0.70 | Yes |
| SCI + Tree + log(Area) | Arachnida – Syrphidae | -0.24 | -0.65 | 0.25 | Yes |
| SCI + Tree + log(Area) | Aves – Arachnida | -0.06 | -0.51 | 0.42 | Yes |
| SCI + Tree + log(Area) | Aves – Apoidea | 0.12 | -0.37 | 0.57 | Yes |
| SCI + Tree + log(Area) | Aves – Mammalia | -0.31 | -0.70 | 0.18 | Yes |
| SCI + Tree + log(Area) | Aves – Syrphidae | -0.40 | -0.76 | 0.08 | Yes |
| SCI + Tree + log(Area) | Mammalia – Arachnida | -0.24 | -0.66 | 0.26 | Yes |
| SCI + Tree + log(Area) | Mammalia – Apoidea | 0.02 | -0.46 | 0.49 | Yes |
| SCI + Tree + log(Area) | Mammalia – Syrphidae | 0.17 | -0.33 | 0.60 | Yes |
| SCI + Tree + log(Area) | Syrphidae – Apoidea | -0.16 | -0.59 | 0.34 | Yes |
| SCI only | Arachnida – Apoidea | 0.34 | -0.10 | 0.70 | Yes |
| SCI only | Arachnida – Syrphidae | -0.11 | -0.53 | 0.33 | Yes |
| SCI only | Aves – Arachnida | -0.05 | -0.48 | 0.38 | Yes |
| SCI only | Aves – Apoidea | 0.23 | -0.22 | 0.61 | Yes |
| SCI only | Aves – Mammalia | -0.31 | -0.67 | 0.14 | Yes |
| SCI only | Aves – Syrphidae | -0.40 | -0.74 | 0.04 | Yes |
| SCI only | Mammalia – Arachnida | -0.23 | -0.62 | 0.22 | Yes |
| SCI only | Mammalia – Apoidea | 0.06 | -0.38 | 0.49 | Yes |
| SCI only | Mammalia – Syrphidae | 0.11 | -0.35 | 0.53 | Yes |
| SCI only | Syrphidae – Apoidea | 0.04 | -0.40 | 0.47 | Yes |

**Table S2** Posterior means and 95% credible intervals of residual correlations between arthropod functional groups. Correlations are derived from multivariate Bayesian models using different combinations of forest structural variables (SCI, dominant tree species, and forest area). Values are based on the residual correlation matrix estimated in *brms*.

| **Model** | **Correlation** | **Estimate** | **CI Lower** | **CI Upper** | **Includes zero** |
| --- | --- | --- | --- | --- | --- |
| Null | Detritivores -Herbivores | -0.03 | -0.45 | 0.40 | Yes |
| Null | Detritivores - Non-feeders | 0.26 | -0.18 | 0.63 | Yes |
| Null | Detritivores - Omnivores | 0.11 | -0.33 | 0.52 | Yes |
| Null | Detritivores - Parasites | -0.22 | -0.60 | 0.22 | Yes |
| Null | Detritivores - Pollinators | -0.05 | -0.47 | 0.38 | Yes |
| Null | Detritivores - Predators | -0.10 | -0.50 | 0.34 | Yes |
| Null | Herbivores - Non-feeders | -0.10 | -0.51 | 0.34 | Yes |
| Null | Herbivores - Omnivores | -0.14 | -0.54 | 0.30 | Yes |
| Null | Herbivores - Parasites | -0.02 | -0.44 | 0.41 | Yes |
| Null | Herbivores - Pollinators | -0.07 | -0.48 | 0.37 | Yes |
| Null | Herbivores - Predators | -0.21 | -0.60 | 0.24 | Yes |
| Null | Non-feeders - Omnivores | 0.21 | -0.23 | 0.59 | Yes |
| Null | Non-feeders - Parasites | 0.15 | -0.28 | 0.54 | Yes |
| Null | Non-feeders - Pollinators | 0.01 | -0.41 | 0.43 | Yes |
| Null | Non-feeders - Predators | 0.14 | -0.30 | 0.54 | Yes |
| Null | Omnivores - Parasites | 0.02 | -0.41 | 0.43 | Yes |
| Null | Omnivores - Pollinators | -0.01 | -0.44 | 0.42 | Yes |
| Null | Omnivores - Predators | -0.04 | -0.46 | 0.40 | Yes |
| Null | Parasites - Pollinators | 0.32 | -0.13 | 0.68 | Yes |
| Null | Parasites - Predators | -0.04 | -0.46 | 0.38 | Yes |
| Null | Pollinators - Predators | -0.11 | -0.52 | 0.33 | Yes |
| SCI only | Detritivores -Herbivores | -0.03 | -0.46 | 0.41 | Yes |
| SCI only | Detritivores - Non-feeders | 0.28 | -0.17 | 0.66 | Yes |
| SCI only | Detritivores - Omnivores | 0.11 | -0.35 | 0.52 | Yes |
| SCI only | Detritivores - Parasites | -0.20 | -0.60 | 0.25 | Yes |
| SCI only | Detritivores - Pollinators | -0.02 | -0.45 | 0.42 | Yes |
| SCI only | Detritivores - Predators | -0.09 | -0.51 | 0.35 | Yes |
| SCI only | Herbivores - Non-feeders | -0.10 | -0.52 | 0.34 | Yes |
| SCI only | Herbivores - Omnivores | -0.13 | -0.55 | 0.32 | Yes |
| SCI only | Herbivores - Parasites | -0.03 | -0.46 | 0.41 | Yes |
| SCI only | Herbivores - Pollinators | -0.09 | -0.51 | 0.36 | Yes |
| SCI only | Herbivores - Predators | -0.20 | -0.60 | 0.25 | Yes |
| SCI only | Non-feeders - Omnivores | 0.22 | -0.23 | 0.61 | Yes |
| SCI only | Non-feeders - Parasites | 0.09 | -0.35 | 0.51 | Yes |
| SCI only | Non-feeders - Pollinators | -0.06 | -0.48 | 0.38 | Yes |
| SCI only | Non-feeders - Predators | 0.13 | -0.31 | 0.54 | Yes |
| SCI only | Omnivores - Parasites | 0.03 | -0.41 | 0.46 | Yes |
| SCI only | Omnivores - Pollinators | 0.01 | -0.43 | 0.45 | Yes |
| SCI only | Omnivores - Predators | -0.03 | -0.47 | 0.41 | Yes |
| SCI only | Parasites - Pollinators | 0.23 | -0.23 | 0.62 | Yes |
| SCI only | Parasites - Predators | -0.06 | -0.48 | 0.38 | Yes |
| SCI only | Pollinators - Predators | -0.13 | -0.55 | 0.32 | Yes |
| SCI + Tree + log(Area) | Detritivores -Herbivores | 0.00 | -0.46 | 0.46 | Yes |
| SCI + Tree + log(Area) | Detritivores - Non-feeders | 0.31 | -0.15 | 0.69 | Yes |
| SCI + Tree + log(Area) | Detritivores - Omnivores | -0.21 | -0.62 | 0.25 | Yes |
| SCI + Tree + log(Area) | Detritivores - Parasites | 0.05 | -0.41 | 0.50 | Yes |
| SCI + Tree + log(Area) | Detritivores - Pollinators | -0.02 | -0.48 | 0.45 | Yes |
| SCI + Tree + log(Area) | Detritivores - Predators | -0.21 | -0.63 | 0.26 | Yes |
| SCI + Tree + log(Area) | Herbivores - Non-feeders | -0.08 | -0.53 | 0.39 | Yes |
| SCI + Tree + log(Area) | Herbivores - Omnivores | -0.12 | -0.55 | 0.35 | Yes |
| SCI + Tree + log(Area) | Herbivores - Parasites | -0.04 | -0.49 | 0.43 | Yes |
| SCI + Tree + log(Area) | Herbivores - Pollinators | -0.06 | -0.53 | 0.43 | Yes |
| SCI + Tree + log(Area) | Herbivores - Predators | -0.19 | -0.62 | 0.29 | Yes |
| SCI + Tree + log(Area) | Non-feeders - Omnivores | 0.26 | -0.20 | 0.65 | Yes |
| SCI + Tree + log(Area) | Non-feeders - Parasites | 0.13 | -0.35 | 0.55 | Yes |
| SCI + Tree + log(Area) | Non-feeders - Pollinators | -0.06 | -0.50 | 0.41 | Yes |
| SCI + Tree + log(Area) | Non-feeders - Predators | 0.28 | -0.19 | 0.67 | Yes |
| SCI + Tree + log(Area) | Omnivores - Parasites | 0.48 | 0.02 | 0.80 | No |
| SCI + Tree + log(Area) | Omnivores - Pollinators | 0.06 | -0.41 | 0.50 | Yes |
| SCI + Tree + log(Area) | Omnivores - Predators | -0.16 | -0.57 | 0.30 | Yes |
| SCI + Tree + log(Area) | Parasites - Pollinators | 0.14 | -0.34 | 0.58 | Yes |
| SCI + Tree + log(Area) | Parasites - Predators | 0.00 | -0.45 | 0.45 | Yes |
| SCI + Tree + log(Area) | Pollinators - Predators | -0.12 | -0.56 | 0.36 | Yes |
| SCI + Tree + SCI:log(Area) | Detritivores -Herbivores | 0.01 | -0.46 | 0.48 | Yes |
| SCI + Tree + SCI:log(Area) | Detritivores - Non-feeders | 0.32 | -0.17 | 0.71 | Yes |
| SCI + Tree + SCI:log(Area) | Detritivores - Omnivores | -0.21 | -0.62 | 0.26 | Yes |
| SCI + Tree + SCI:log(Area) | Detritivores - Parasites | 0.06 | -0.42 | 0.51 | Yes |
| SCI + Tree + SCI:log(Area) | Detritivores - Pollinators | -0.03 | -0.50 | 0.45 | Yes |
| SCI + Tree + SCI:log(Area) | Detritivores - Predators | -0.21 | -0.63 | 0.27 | Yes |
| SCI + Tree + SCI:log(Area) | Herbivores - Non-feeders | -0.12 | -0.57 | 0.37 | Yes |
| SCI + Tree + SCI:log(Area) | Herbivores - Omnivores | -0.16 | -0.59 | 0.31 | Yes |
| SCI + Tree + SCI:log(Area) | Herbivores - Parasites | -0.04 | -0.50 | 0.45 | Yes |
| SCI + Tree + SCI:log(Area) | Herbivores - Pollinators | 0.02 | -0.47 | 0.51 | Yes |
| SCI + Tree + SCI:log(Area) | Herbivores - Predators | -0.28 | -0.68 | 0.22 | Yes |
| SCI + Tree + SCI:log(Area) | Non-feeders - Omnivores | 0.24 | -0.23 | 0.64 | Yes |
| SCI + Tree + SCI:log(Area) | Non-feeders - Parasites | 0.12 | -0.37 | 0.56 | Yes |
| SCI + Tree + SCI:log(Area) | Non-feeders - Pollinators | -0.01 | -0.48 | 0.47 | Yes |
| SCI + Tree + SCI:log(Area) | Non-feeders - Predators | 0.24 | -0.24 | 0.65 | Yes |
| SCI + Tree + SCI:log(Area) | Omnivores - Parasites | 0.46 | -0.02 | 0.80 | Yes |
| SCI + Tree + SCI:log(Area) | Omnivores - Pollinators | 0.10 | -0.38 | 0.54 | Yes |
| SCI + Tree + SCI:log(Area) | Omnivores - Predators | -0.20 | -0.61 | 0.27 | Yes |
| SCI + Tree + SCI:log(Area) | Parasites - Pollinators | 0.14 | -0.36 | 0.59 | Yes |
| SCI + Tree + SCI:log(Area) | Parasites - Predators | 0.00 | -0.47 | 0.47 | Yes |
| SCI + Tree + SCI:log(Area) | Pollinators - Predators | -0.04 | -0.51 | 0.44 | Yes |
